# Altered hypothalamic microstructure in human obesity

**DOI:** 10.1101/593004

**Authors:** K. Thomas, F. Beyer, G. Lewe, R. Zhang, S. Schindler, P. Schönknecht, M. Stumvoll, A. Villringer, A.V. Witte

## Abstract

Obesity is a multifactorial disorder driven by sustained energy imbalance. The hypothalamus is an important regulator of energy homeostasis and therefore likely involved in obesity pathophysiology. Animal studies suggest that obesity-related diets induce structural changes in the hypothalamus through inflammation-like processes. Whether this translates to humans is however largely unknown. Therefore, we aimed to assess obesity-related differences in hypothalamic macro- and microstructure based on a multimodal approach using T1-weighted and diffusion-weighted magnetic resonance imaging (MRI) acquired at 3 Tesla in a large well-characterized sample of the Leipzig Research Center for Civilization Diseases (LIFE) cohort (*n*_1_ = 338, 48% females, age 21-78 years, BMI 18-43 kg/m^2^). We found that higher body mass index (BMI) selectively predicted higher mean proton diffusivity (MD) within the hypothalamus, indicative of compromised microstructure in the underlying tissue. Results were independent from confounders and confirmed in another independent sample (n_2_ = 236). In addition, while hypothalamic volume was not associated with obesity, we identified a sexual dimorphism and larger hypothalamic volumes in the left compared to the right hemisphere. Using two large samples of the general population, we showed that a higher BMI specifically relates to altered microstructure in the hypothalamus, independent from confounders such as age, sex and obesity-associated co-morbidities. This points to persisting microstructural changes in a key regulatory area of energy homeostasis occurring with excessive weight. These findings may help to better understand the pathomechanisms of obesity and other eating-related disorders.

## Introduction

Obesity is associated with dysfunctions in central homeostatic regulation, which might also play a pivotal role in its pathogenesis^1,2^. Energy homeostasis (i.e. the balance between food intake and energy expenditure) depends on signaling pathways in the hypothalamus, a small diencephalic brain region comprised of different sub-nuclei^3,4^ Here, distinct subpopulations of neurons integrate circulating hormones that signal satiety (e.g. leptin, insulin) and hunger (e.g. ghrelin)^5^.

Animal models support the hypothesis that a high-fat diet (HFD) triggers an inflammation-like response in the hypothalamus, which in turn impairs the sensing of anorexigenic signals, thereby contributing to continuous food intake and weight gain^6,7^ For example, rodents fed a HFD showed increasing levels of proinflammatory cytokines such as interleukin-6 (IL-6) and tumor necrosis factor alpha (TNF*α*) in the hypothalamus^8^, even prior to substantial weight gain^9^ This immunologic response was also accompanied by a rapid accumulation of microglia and recruitment of astrocytes^7,10^ Additionally, hypothalamic neurons showed signs of toxic stress and underwent apoptosis after HFD^11^. While some studies reported that this inflammation-like response declined after several days of overnutrition, suggesting a compensatory mechanism to prevent neurons from damage^12^, others showed that gliosis and astrocytosis reoccurred after several weeks, pointing to prolonged changes in hypothalamic tissue and microstructural properties in obese animals^9^.

Whether these neurobiological alterations shown in animal models of obesity also contribute to the pathophysiology of obesity in humans is however largely unknown. A post mortem analysis of obese and non-obese individuals reported that a higher BMI correlated with alterations in hypothalamic glia cells, which exhibited increased levels of dystrophy according to histological stainings^12^. Studies using *in vivo* magnetic resonance imaging (MRI) linked volumetric changes in the hypothalamus to altered eating behavior within neurodegenerative and psychiatric disorders such as frontotemporal dementia or schizophrenia^13–15^ Two studies provided initial evidence for changes in hypothalamus T2-weighted magnetic resonance imaging (MRI) signals in relation to obesity: Thaler et al. showed increased signal ratio in a circular region-of-interest (ROI) in the left hypothalamus referenced to an amygdala-ROI in 12 obese compared to 11 non-obese participants^9^. Another study including 67 participants reported higher T2-relaxation times in obesity within a ROI in the left mediobasal hypothalamus, and both studies proposed these measures as a marker of hypothalamic gliosis in diet-induced obesity^9,16^ However, sample sizes were small and applying fixed ROIs could be misleading due to partial volume effects and the heterogenous appearance of the hypothalamus. In addition, the direction of effects was partly contradictory and a limited resolution and multiple sources of image artifacts limit interpretability^14,15,17–19^.

In sum, animal experiments and first, but not all, human studies support the hypothesis that central homeostatic changes reflected in compromised (micro)structure of the hypothalamus are present in obesity. However, methodology in the human studies remained unconvincing so far^18,20–22^. We therefore applied advanced voxel-wise MRI techniques^23,24^ to determine whether larger hypothalamic volume and higher hypothalamic mean diffusivity (MD), derived by diffusion tensor imaging (DTI) and commonly interpreted as less intact cellular microstructure^25,26^, are positively associated with obesity measured using BMI in a well-characterized large population-based sample. We also explored whether hypothalamic MD was linked to higher visceral adipose tissue volume (VAT), given the elevated inflammatory risk profile of this body fat depot^27^ We additionally implemented a multi-atlas based label segmentation to validate our results in another independent sample.

## Results

### Hypothalamic volume

In a sample of 338 participants (48% females, aged 21-78 years, BMI range of 18-43 kg/m2), we delineated the left and right hypothalamus on T1-weighted anatomical MRI using a state-of-the-art semi-automated segmentation algorithm resulting in individual hypothalamic masks at the voxel-level (**Fig. 1A**, see **Methods** for details).

**Figure 1:**
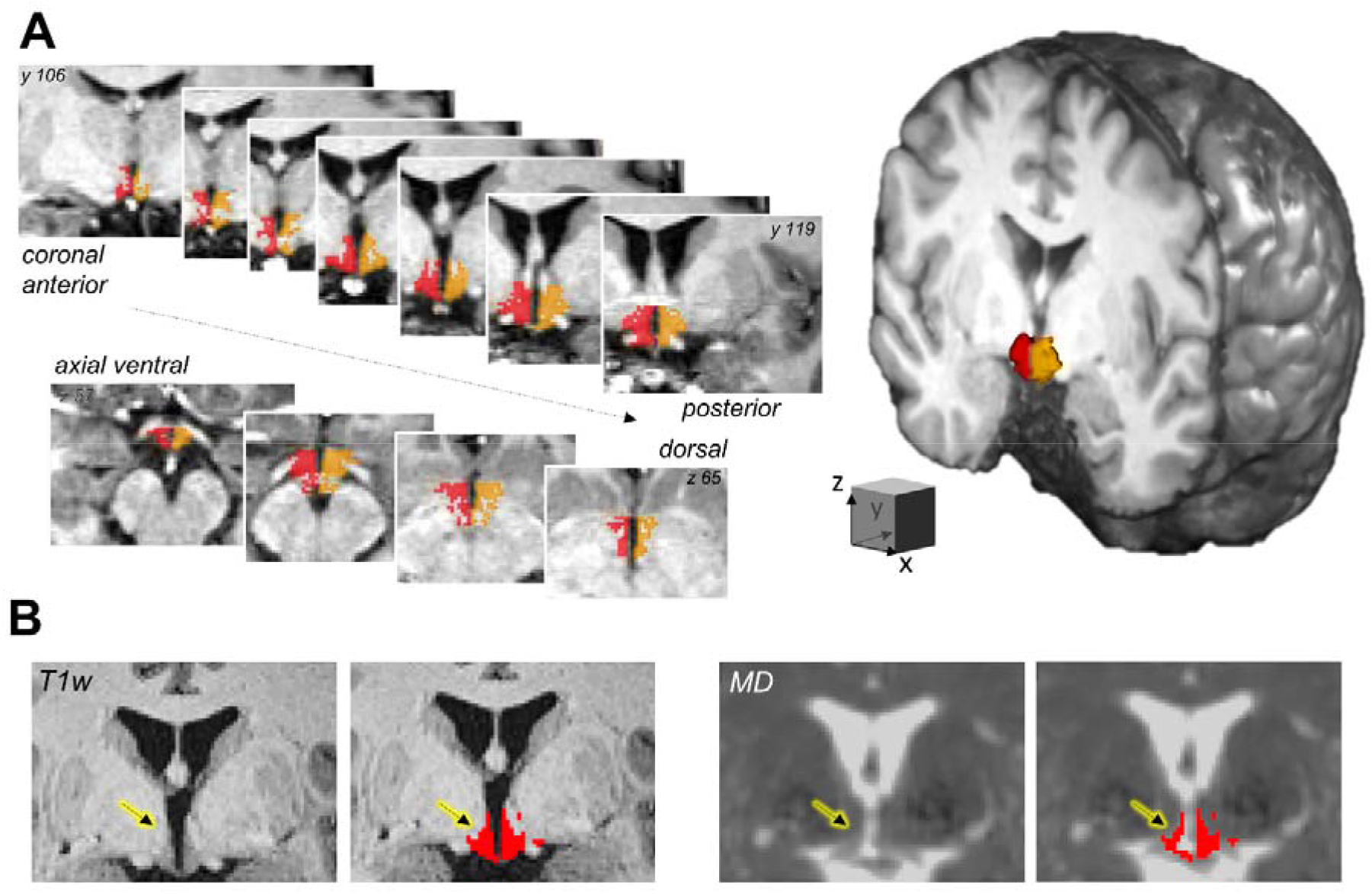
The hypothalamus on multimodal MRI. **A**: The bilateral hypothalamus (right: red, left: orange) of a representative participant according to semi-automated segmentation on anatomical images. **B**: Coregistration of the T1-weighted (T1w)-derived hypothalamus mask to the mean diffusivity (MD) image derived by diffusion-weighted imaging. Note the sparing of hypothalamus voxels which are affected by partial volume effects on the MD image (arrows). Images are shown in radiological convention.

On average, men showed 12.8% larger head-size adjusted whole hypothalamic volumes than women (*n*_1_ = 338; 48% females, aged 21-78 years, BMI range of 18-43 kg/m^2^; **Fig. 2**). The difference was statistically significant (β=-0.18, p < 0.001) according to a multiple regression model which explained 21.8% of the variance (F_3,333_ = 26.9, p < 0.001) and controlled for potential effects of age (no significant contribution, p = 0.96), and rater (β_0,1_=-0.56, p < 0.001, β_0,2_=-0.33, p=0.001). Adding BMI as additional predictor to the model did not improve the model fit (p = 0.58) indicating that BMI was not associated with hypothalamic volume.

**Figure 2:**
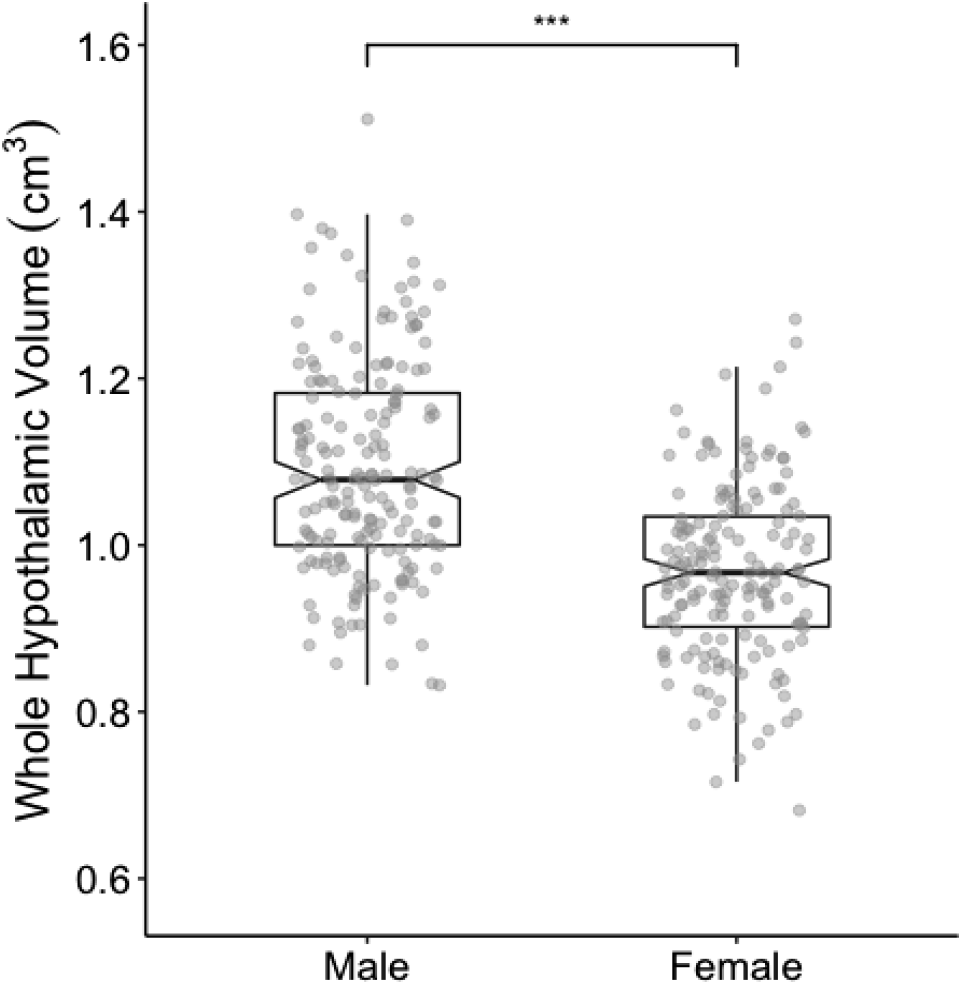
Sex differences in hypothalamic volume. Analysis of hypothalamic volume reveals bigger values for male than for female participants (p < 0.001).

When investigating the hemispheres separately, we observed higher volume for the left than for the right hypothalamus, an effect which was slightly less pronounced in women and independent of age and rater (linear mixed effect model, side: β=-40.9; sex: β=-26.5; side-by-sex interaction: β=1.98; p = 0.048; rater: β_0,1_=-52.6/ β_0,2_=-39.3, p<0.001; age:p = 0.48).

### Obesity and hypothalamic microstructure

Next, we examined average MD within the individual’s hypothalamus using DTI as a sensitive measure of microstructural properties^25,28^. A carefully designed processing pipeline ensured that DTI-related distortions adjacent to the hypothalamus region did not bias hypothalamic MD estimates (**Fig. 1B**, see **Methods** for details).

According to linear regression, BMI significantly predicted hypothalamic MD (β = 14, p = 0.008), showing that higher BMI was related to higher MD (**Fig. 3A**). The regression model (F_4,306_ = 24.5, p < .001, R^2^ = 0.23) adjusted for potential effects of sex (β = −0.19, p < 0.001), age (β = 0.38, p < 0.001), and rater (n.s., p = 0.67). Men had larger MD than women and higher age was linked to higher MD. Adding BMI as predictor explained 1.5% more variance in hypothalamic MD than a model without BMI (F_1,306_ = 7.1, p = 0.008). To test the specificity of our findings, we added MD within the hippocampus as another heterogenous subcortical structure to the model, which did not attenuate the predictive association of BMI and hypothalamic MD. The same was true when adding the volume of the 3rd ventricle or the hypothalamic volume as covariate to the model, to account for partial volume effects.

**Figure 3:**
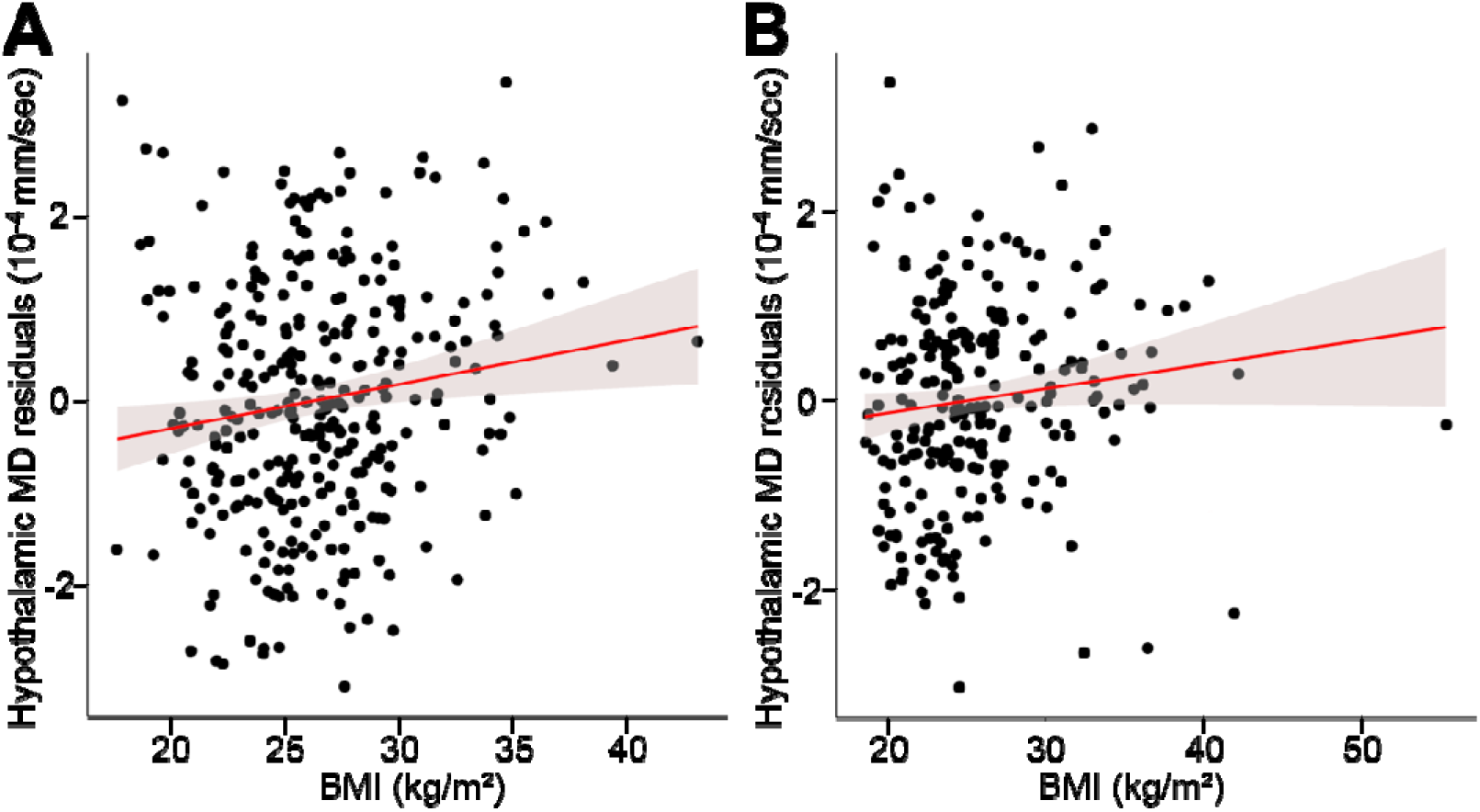
Obesity and hypothalamic microstructure. Higher body mass index (BMI) significantly predicts higher hypothalamic mean diffusivity (MD), commonly interpreted as less intact cellular microstructure, in a first (A, n_1_ = 311, comparison to age, sex-corrected model, F_1,306_ = 7.1, p = 0.008) and a second independent sample (B, n_2_ = 236, comparison to age, sex-corrected model, F_1,232_ = 4.2, p = 0.041). Line indicates regression fit with 95% confidence interval.

### Confirmatory analysis

To validate our findings in an independent sample, we developed a novel multi-label fusion atlas based on the initial segmentations that automatically generates individual hypothalamic segmentations (**Fig. 4**; see **Methods** for details). Using this atlas-approach in a second group of 236 participants confirmed a significant association between higher BMI and higher hypothalamus MD in similar magnitude (β = 0.14, p = 0. 04, **Fig. 3B**; regression model: F_3,232_ = 15.5, p < .001, R^2^ = 0.41), adjusted for age (β = 0.37, p < 0.001) and sex (β = −0.12, p = 0.04). Changes in F-values confirmed that adding BMI increased the explained variance of hypothalamic MD significantly by 1.5% (F_1,232_ = 4.2 p = 0.04). Similar to the initial sample, when adding hippocampal MD and ventricular volume to the model, BMI remained a significant predictor of hypothalamic MD. Furthermore, consideration of obesity-associated biomarkers (systolic blood pressure and HOMA-IR) as possible confounders did also not attenuate the positive association between BMI and hypothalamic MD. We report good to excellent reliability (ICC > 0.87) between the semi-automated and the fully-automated segmentation procedures for hypothalamus MD.

**Figure 4:**
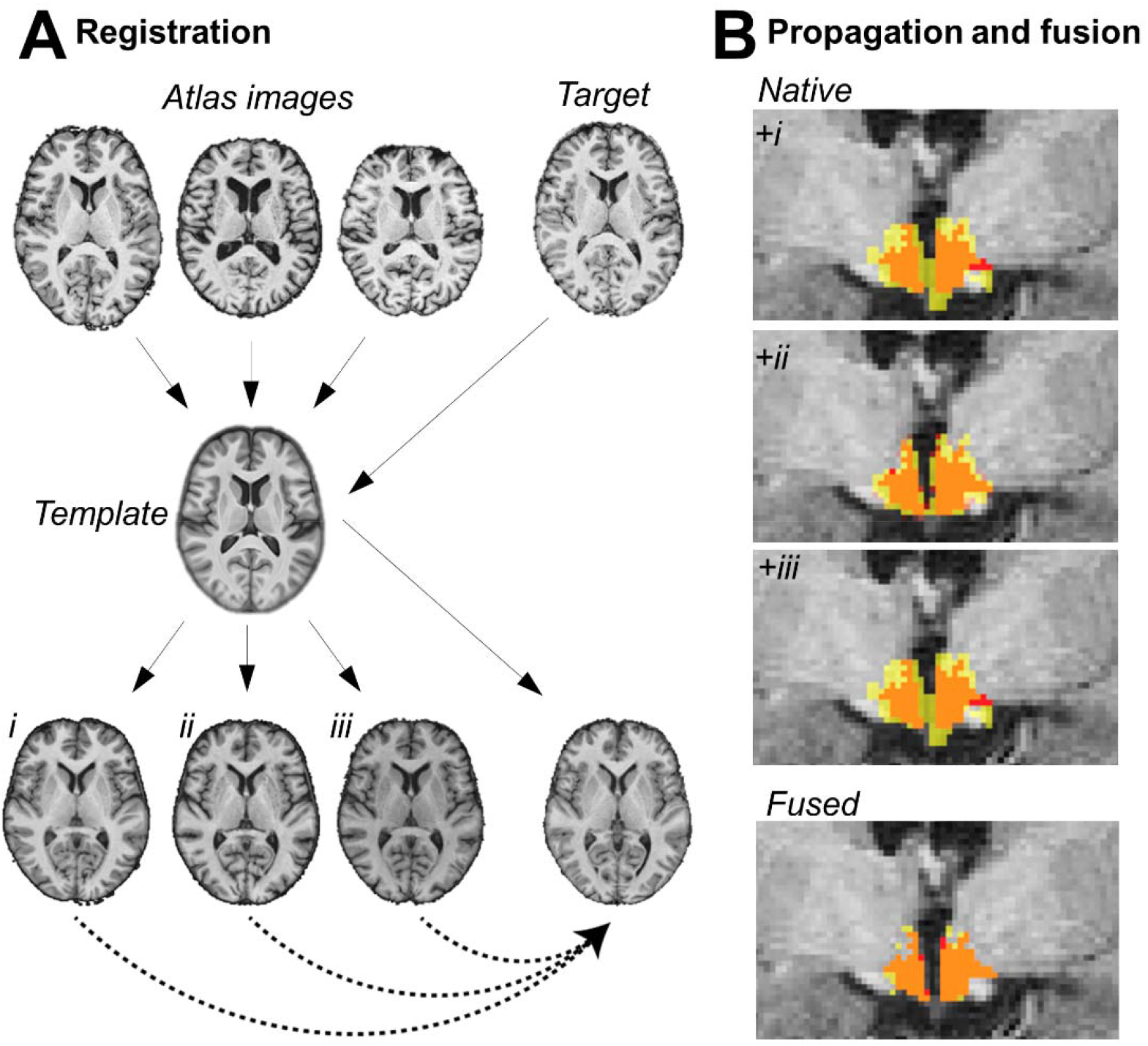
Multi-atlas fusion segmentation for automated hypothalamus segmentation. **A**: In the registration step both atlas and target images were nonlinearly registered to a template image. In this common space another non-linear registration of atlas images to the target image was performed. **B**: In the label propagation step all transformations were concatenated and the atlas hypothalami were brought into the native space of the target image (upper images, yellow: propagated label, red: manual label, orange: overlap). Fusion of the region of interest was performed using STEPS (lower image, yellow: fused label, red: manual label, orange: overlap see text for details).

### Exploratory analysis of visceral fat

To explore whether visceral obesity explained additional variance in hypothalamic MD, we added log-transformed height-corrected VAT as an additional predictor into the regression model (F_5,300_ = 19.8, p < .001, R^2^ = 0.24). VAT was estimated from T1-weighted abdominal MRI in a subset of the initial sample (n = 306, see **Methods** for details). Controlling for the impact of age (β = 0.37, p < 0.001), sex (β = −0.17, p < 0.001), rater (p = 0.658) and BMI (p = 0.132), we did not find significant associations for VAT (F_1,300_ = 0.5, p = 0.5) and average hypothalamic MD.

## Discussion

Using multimodal neuroimaging in two large samples of healthy adults, we showed that higher BMI is associated with higher proton diffusivity in the hypothalamus, indicating hypothalamic microstructural alterations in obesity. In parallel, while men had higher hypothalamus volumes than women and the volume of the left hemispheric hypothalamus was larger than the right, BMI was not associated with hypothalamic volume.

### Hypothalamic microstructure and obesity

Our findings provide evidence that higher BMI is associated with compromised microstructure in the hypothalamus. While the effect size is to be considered small, explaining 1.5% of the variance in hypothalamus MD, we confirmed our findings in another large independent sample. Our results are in line with and extend previous animal and human studies reporting obesity-related alterations in hypothalamic microstructure assessed with T2-weighted imaging, though previous human studies were based on limited sample sizes, suffered from two-dimensional assessments of the hypothalamus and have used less established markers of microstructure^9,20,22,29,30^ In contrast, DTI-derived MD in grey matter regions, as used in the current study, reflects the amount, density or integrity of neuronal membranes, dendrites, axons, or glial compartments, that restrict water diffusion in the tissue in both animals and humans^25,26,29^ Previous work showed that higher MD for example in the hippocampus correlated with poorer memory function^28^ This might indicate that obesity-associated higher MD in the hypothalamus goes along with microstructural changes that could lead to dysfunctional outcomes. We also found higher values of hypothalamic MD in men than in women as well as an age-related increase in MD. The latter is supported by a broad range of studies that consistently found positive associations between diffusion metrics (such as MD and FA) and age in various GM structures, often in line with worse cognitive performance^31,32^.

Yet, despite of being able to detect alterations on a cellular level, DTI metrics such as MD suffer from non-specificity and are confounded by tissue geometry. Accordingly, MD has been linked to various neurological disorders as well as to unspecific cerebral abnormalities such as edema, necrosis, demyelination or augmented cellularity^25^. Therefore, various underlying mechanisms might explain the obesity-associated increases in hypothalamic MD in our study.

First, as discussed in the concept of hypothalamic inflammation, changes in MD might be attributed to a sustained gliosis as a consequence of diet-induced obesity. This is supported by findings in mice showing that microgliosis and astrocytosis returned permanently in mice fed a HFD, although temporarily subsiding^9^ In addition, another study suggested microglial responses due to ongoing malnutrition in humans as they also detected signs of gliosis and microglial dystrophy in human hypothalamus assessed by post mortem stereology^12^.

Second, hypothalamic inflammation in mice is also linked to a loss of hypothalamic neurons that underwent apoptosis as a consequence of the HFD^11^. Therefore, the observed diffusion alteration might also be due to an enhanced amount of extracellular fluid that is accompanied by the neuronal loss or the neuroinflammation in general^33^.

Another possible explanation for the increase in MD addresses vessel integrity, as it has been shown that HFD triggers hypothalamic angiopathy in mice with increased vessel density and length^34^ Currently, new approaches are underway that aim to disentangle the changes in diffusion metrics driven by blood perfusion originating from the extracellular space^35^

Taken together, MD was positively associated with BMI in two large samples. While this suggests small, but reliable alterations in the hypothalamic microstructure in obese humans, the underlying histological mechanisms remain elusive. We encourage future studies to link our neuroimaging findings with advanced analysis at the cellular level (e.g. post-mortem stereology) to further explore the underlying mechanisms.

### Hypothalamic volume

Our voxel-wise estimations of hypothalamic volume in a total of 338 participants, which is the largest sample of hypothalamic volumes obtained by semi-automated segmentation to date, adds to previous reports that whole hypothalamic volume assessed by MRI techniques is around 1 cm^3^ [49]. Reliability analysis revealed acceptable to excellent intra-rater and inter-rater reliabilities of hypothalamus volume and spatial overlap of resulting masks using this method. This highlights the sensitivity and specificity of our procedure and compares to previous high-quality segmentation protocols implemented in smaller sample sizes^13,19,36^.

We also found that hypothalamic volumes were higher for males compared to females, irrespective of head size, age and BMI. This finding might be attributable to known sex differences in metabolic dysregulation^37^ and the neuroendocrine regulation system^38^ Furthermore, we found a significant left-right asymmetry in hypothalamic volume with higher volumes for the left than for the right hypothalamus, which is in line with a previous publication that described a trend in the same direction in a sample of 84 subjects^24^ Along these lines, some hypothalamic functions have been described as lateralized to the left^39^ Recent studies also suggest the hypothalamus to be involved in a lateralized brain circuit that mediates feeding behavior and homeostatic regulation^40^. Future studies need to explore whether these processes might also contribute to volumetric asymmetry in hypothalamic volume.

We did not find a significant relationship between hypothalamic volume and BMI, controlling for the impact of age, sex and the different raters. Although a wide range of literature demonstrate that higher BMI is associated with lower GM volumes in various brain regions^41^, evidence for significant changes in hypothalamic volumes associated with obesity is less observed^42,43^ Nevertheless, BMI has been shown to be related to functional alterations in several brain circuits that involve the hypothalamus^44^. Interestingly, while age-related atrophy in various subcortical structures is commonly observed^45^, age was not related to hypothalamic volume in the present cohort.

### Limitations and strengths

Some limitations need to be taken into consideration. As our dataset is crosssectional, we cannot infer causality. Altered hypothalamic microstructure might be attributable to both, prerequisite or consequence, of obesity. Furthermore, even if referring to established concepts such as hypothalamic inflammation, knowledge about the temporal dynamics of this inflammatory process is scarce or inconsistent^12^. Also, hypothalamus physiological function is not restricted to energy metabolism and homeostasis, and we were not able to dissect the hypothalamus in its sub-nuclei. Thus, we cannot rule out whether the arcuate nucleus, although relatively large, and/or other hypothalamus subnuclei, serving as main hubs in the control of fluid balance, circadian rhythms or thermoregulation^46^, contributed to the average hypothalamic MD signal. In addition, the usage of BMI to characterize obesity might be too simplistic^27^ However, our results incorporating MRI-based measures of VAT, indicative of visceral obesity, strongly indicate that VAT did not improve the model fit with regard to microstructural changes within the hypothalamus. Arguments for the robustness and specificity of our findings stem from covariate adjustments for age, sex and other potential confounders. Particularly, considering hippocampal MD, HOMA-IR and systolic blood pressure in our statistical analysis did not attenuate the association between obesity and hypothalamic MD in our validation sample. Reliability analyses indicated good to excellent fits between the MD-methods used in the two samples. Further strengths of our study include the large, well-characterized population-based sample size, a thorough methodological design combining a semiautomated segmentation algorithm with sensitive DTI metrics, along with confirmation analysis in an independent sample.

## Conclusion

Using a novel multimodal MRI approach in two large samples of healthy adults of the general population, we were able to demonstrate that a higher BMI specifically relates to higher MD in the hypothalamus, independent from confounders such as age, sex and obesity-associated co-morbidities. This finding thus points to persisting microstructural alterations in a key regulatory area of energy homeostasis occurring with excessive weight. The underlying mechanisms might include inflammatory activity, neuronal degeneration or angiopathy in the hypothalamus due to obesity-related overnutrition and metabolic alterations. Future studies need to test the functional relevance of these microstructural changes, and if interventions aiming to reduce obesity can effectively reverse the observed changes in hypothalamic MD.

## Material and Methods

### Participants

Participants were recruited randomly as part of the MRI-subsample within the “Health Study for the Leipzig Research Centre for Civilization Diseases” (LIFE-Adult) study^47^ The study was approved by the Ethics Committee of the University of Leipzig and all participants gave informed written consent. In total, 2637 adults received brain MRI. We selected participants without history of stroke, cancer, epilepsy, multiple sclerosis and Parkinson’s disease, neuroradiological findings of brain pathology or intake of centrally active medication (n = 2095, for a flowchart, see **Fig. 5**). Further, only a well-characterized subgroup with abdominal MRI to assess visceral adipose tissue (VAT) was considered (n = 993). Out of these, two raters segmented 166 and 152 participants, respectively. For test-retest and interrater-reliability both raters additionally segmented 20 participants twice. In total, bilateral hypothalami were segmented in n = 338 participants (*n*_1_, for demographic characteristics, see **Table 1**). Twenty-seven participants had to be removed from diffusion-weighted image analysis due to incomplete or deficient imaging data. For confirmatory analyses, we additionally examined another n = 236 of the pool of participants with additional abdominal MRI using multi-atlas fusion segmentation (*n*_2_, see Fig. 5 and below for details).

**Figure 5:**
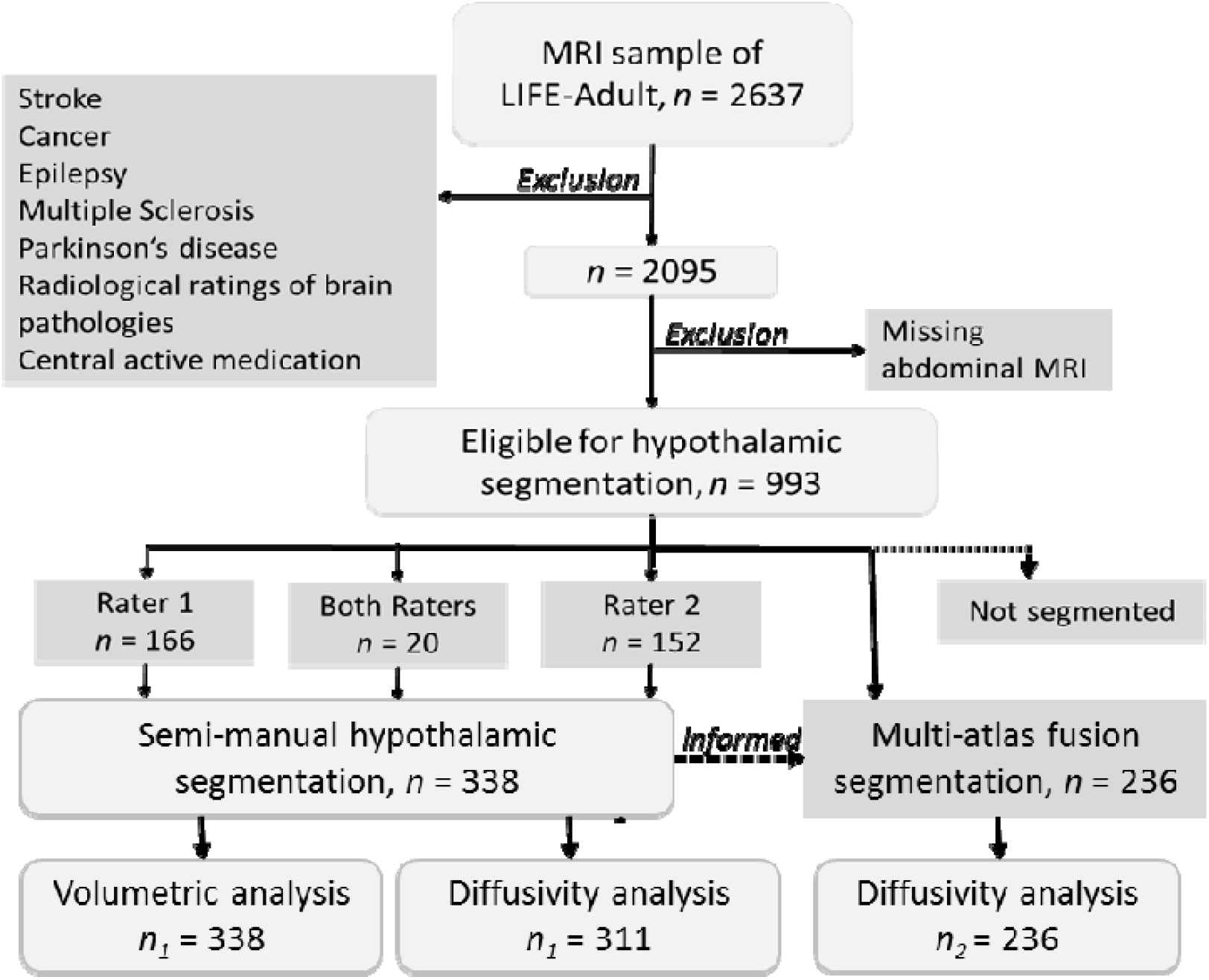
Flowchart of the study illustrating the exclusion criteria, the subsample sizes and the different approaches of data analysis.

**Table 1:**
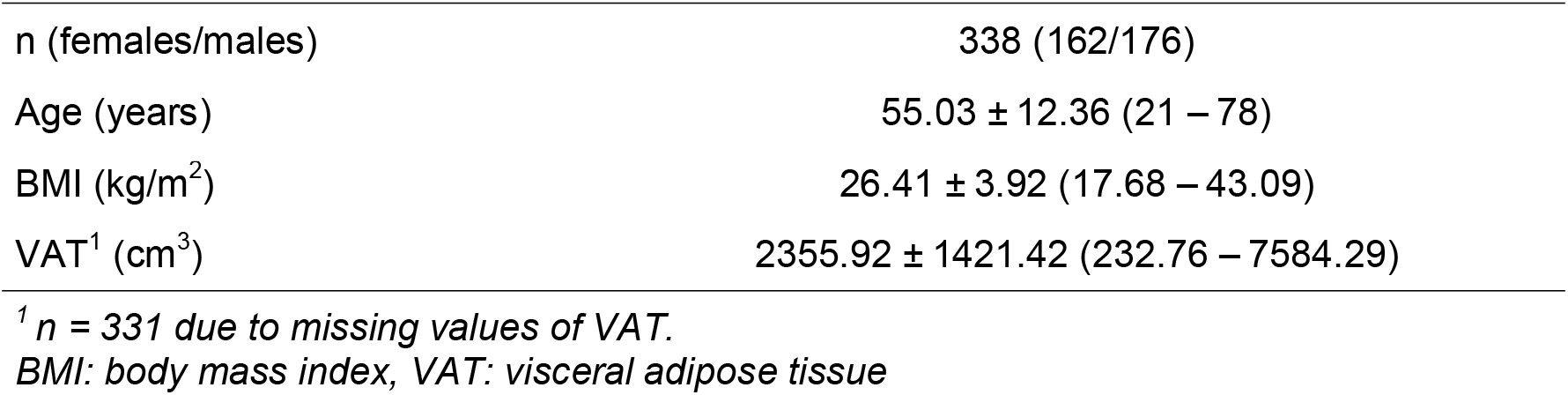
Demographic characteristics of the semi-manual segmentation sample n_1_. Data is given as mean ± standard deviation (SD) and range (minimum – maximum).

### Anthropometry

Body weight was measured with a scale with a precision of 0.01kg and body height was assessed using the means of a stadiometer to the nearest 0.1cm. BMI was calculated as body weight [kg] divided by squared body height [m].

### Obesity-related biomarkers

We collected additional obesity-related biomarkers in a subset of participants. Laboratory indicators of glucose metabolism (glucose and insulin) were obtained after overnight fasting according to standard procedures^47^ and used to calculate insulin resistance with the homeostatic model assessment (HOMA-IR)^48^ Blood pressure was measured with an automatic oscillometric blood pressure monitor (OMRON 705IT, OMRON Medizintechnik Handelsgesellschaft mbH).

### Magnetic Resonance Imaging

Magnetic resonance imaging (MRI) was performed on a 3T Magnetom Verio scanner (Siemens, Erlangen, Germany, equipped with a 32-channel head array coil and syngo MR B17 software).

### Abdominal MRI acquisition and preprocessing

MRI of the abdomen was performed using an axial T1-weighted fast spin-echo technique with the following parameters: repetition time, 520 ms; echo time, 18 ms; 5-mm gap between slicefield of view, 500 mm 375 mm; final voxel size 1.6 1.6 5.0 mm^3^. Beginning 10 cm below the umbilicus, 5 slices were recorded from feet-to-head direction with 5 cm table shift after each acquisition and finishing in the liver region^47^ Using a semi-automated segmentation algorithm implemented in ImageJ (https://imagej.nih.gov/ij/download/), visceral adipose tissue (VAT) was obtained from 20 slices centered around the participant’s umbilicus^49^ For subsequent analysis, the VAT volume was log-transformed and normalized by height.

### Head MRI acquisition and preprocessing

Anatomical MRI was acquired using a T1-weighted Magnetization prepared rapid gradient echo (MPRAGE) pulse sequence with the following parameters: inversion time, 900 ms; repetition time, 2.3 ms; echo time, 2.98 ms; flip angle, 98;; image matrix, 256 176 240; voxel size, 1×1×1 mm^3^.

Preprocessing of the anatomical T1-weighted data included skullstripping and realignment to anterior and posterior commissure in Lipsia (https://www.cbs.mpg.de/institute/software/lipsia/download). Then, tissue segmentation was performed with the default settings using SPM12’s New Segment based on Matlab version 2017b.

Diffusion tensor imaging (DTI) was acquired with a twice-refocused echo planar imaging sequence (EPI) with the following parameters: repetition time, 13800 ms; echo time, 100 ms;; image matrix 128 128; 72 slices; voxel size 1.7 1.7 1.7mm^3^; 60 directions with b-value 1000 s/mm^2^, and 7 volumes with b-value 0s/ mm^2^.

Preprocessing included denoising (MRtrix v3.0) of the raw data removal of gibbs-ringing artifact from all b0 images using the local subvoxel-shift method and outlier replacement using the eddy tool in FSL 5.0.10^50–53^. Subsequently, data was corrected for head motion and linearly coregistered to the T1 image with Lipsia tools. Finally, we applied tensor model fitting and generated mean diffusivity (MD) and fractional anisotropy (FA) images.

### Semi-automated segmentation of the hypothalamus

Based on previously established protocols for 3T MRI data^24^, we performed semiautomated segmentation of the hypothalamus in MeVisLab 4.1. Briefly, a preoptic, an intermediate-superior and -inferior as well as a posterior region of interest (ROI) was manually pre-defined by two raters using the following landmarks: anterior commissure, columna fornicis, interventricular foramen, mamillary bodies, zona incerta and hypothalamic sulcus^24^ Due to some false-positive segmentation results of intraventricular voxels with the original approach, we adapted the medial landmarks according to Mai, Majtanik & Paxinos (2015). Next, grey matter tissue probability masks were overlaid on each ROI and predefined grey matter thresholds were used to define hypothalamus area. Then all slices were combined to generate a three-dimensional volume of the hypothalamus (**Fig. 2A**). Subsequently, each rater checked the results carefully in a triplanar view with regard to plausibility and coherence to the predefined anatomical edges.

To perform intra- and interrater reliability analysis, we selected 20 additional participants that were segmented twice by both raters. We ensured that reliability subjects were comparable to the whole segmentation sample with respect to age, sex and BMI. According to Shrout & Fleiss (1979), intraclass correlation coefficients (ICC) were calculated using model 1,1 and 3,1. We considered an ICC□≥□0.9 excellent, 0.9□>□ICC□≥□0.8 good and 0.8□>□ICC□≥□0.7 acceptable^56^. Additionally, percentage of relative overlap between the two raters was assessed using Dice similarity coefficient (DSC)^57^. An overlap of 70, 80 or 90% (DSC = 0.7, 0.8, 0.9) was regarded acceptable, good and excellent, respectively. All ICC and DSC values showed acceptable to excellent agreements (Supplementary Tab. 1).

The segmentation procedure was conducted separately for left and right hypothalamus and took between 30 and 45 minutes per brain. Hypothalamic volumes were assessed by extracting the number of voxels for each side. Whole hypothalamic volume was calculated by summing up volumes of left and right hypothalamus. As subcortical volumes are trivially linked to total intracranial volume, hypothalamic volume was adjusted using the following formula^58^:

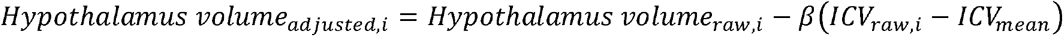

where ICV is the total intracranial volume and *β* is the unstandardized slope of a regression model between ICV and the whole hypothalamic volume across participants. As nonparametric Shapiro-Wilk test indicated a non-normal distribution of the adjusted hypothalamic volumes, we log-transformed volumetric data.

For the statistical analysis, we considered rater as a variable with three levels: rater 1, rater 2 and “rater1/2”. For these 20 reliability subjects, we used the average of the two measurements by the two raters.

### Extraction of the hypothalamus mean diffusivity

Derived by DTI, we used MD as a sensitive measure of microstructural properties^25,28^ Briefly, MD reflects the overall amount of diffusion in a certain voxel, and we averaged this measure in the hypothalamic ROI. A carefully designed processing pipeline ensured that DTI-related distortions adjacent to the hypothalamus region did not bias hypothalamic MD estimates (**Fig. 2B**). FA images of all subjects with hypothalamic volumetry were coregistered to the respective anatomical images with FSL’s FLIRT using 6 degrees of freedom. Then, the registration matrix was used to coregister the MD images to the anatomical space. 24 participants did not receive diffusion weighted imaging or had incomplete data. Furthermore, coregistration failed in 3 subjects, resulting in 311 participants eligible for MD analysis in sample *n*_1_.

Due to its small size, minor shifts or artifacts within the overlay of hypothalamus and the MD mask might be detrimental for analysis, especially for hypothalamic and non-hypothalamic voxels adjacent to the third ventricle (Fig. 2B). In order to avoid that intraventricular voxels were regarded as hypothalamic tissue, further processing was required to distinguish these voxels from those in hypothalamic tissue with regard to MD. Consequently, we derived the average MD in the third ventricle based on the automatic segmentation in FreeSurfer version 5.3.0. Suggesting that grey matter (hypothalamus) MD is smaller than MD in cerebrospinal fluid (third ventricle)^59^, average MD of the whole third ventricle was chosen as a threshold for the hypothalamic MD. Specifically, the MD value of each putative hypothalamic voxel was compared to the average MD of the whole third ventricle. Unless MD of each voxel was higher than the average MD of the third ventricle, this voxel was considered hypothalamic. Results were manually crosschecked. Finally, average MD of all voxels that were likely to be hypothalamic tissue was extracted.

### Statistical analysis of hypothalamic volume and diffusivity

R version 3.2.3 was used to perform statistical analysis.

BMI-related differences in whole hypothalamic volume and MD were assessed by two groups of regression models. For both hypothalamic volume and MD, we compared the null model (including age, sex and rater as predictors) against a regression model including BMI as an additional predictor. The difference between the model was assessed using a F-test and a p-value < .05 was regarded as statistically significant. To test the specificity of the finding and exclude confounding of ventricular volume, we additionally tested a model including the MD of the hippocampus and the ventricular volume as predictors^28^

Hemispheric and sex differences of hypothalamic volume were evaluated in a linear mixed model which included a side-by-sex interaction, rater and age as predictors and subject as a random factor. We report β estimates and p-values based on likelihood ratio tests-based for the fixed main and interaction effects.

### Confirmatory analyses

#### Multi-atlas fusion segmentation

We aimed to confirm the above described MD-analyses in another independent sample. Therefore, we implemented a fully automated multi-label fusion hypothalamus segmentation procedure (**Fig. 3**). First, we created a study-specific template. We used n = 150 randomly selected participants with manual segmentations of the hypothalamus out of sample *n*_1_. This sub-sample did not differ from the final sample (*n*_1_ = 338) in age, sex, BMI or rater distribution (all p > 0.05) (Supplementary Table 4).

To create the template, we applied the function buildtemplateparallel.sh implemented in ANTS version 2.2.0^60^ For more details on the code see publicly available scripts (https://edmond.mpdl.mpg.de/imeji/collection/wLm6DPKVY7_yIzyz). We then implemented a multi-atlas label fusion based on an intermediate template in nipype (for details, see Supplementary information)^61–63^

Finally, we extracted the volume of the resulting hypothalamus segmentation and the ventricle-thresholded average MD values. We validated this approach in two samples.

First, we performed the multi-label fusion segmentation for each of the 44 participants from the template sample. We compared estimated hypothalamic volumes and MD with the values derived from the manual segmentation using ICC (model 3,1) and DSC. In this sample, three participants could not be included for the analysis of MD due to deficient DTI preprocessing.

In the second validation, we aimed to test whether the automated segmentation would perform equally well in participants who were not included into the template. Therefore, we randomly selected 24 participants with manual segmentations who were not part of the n = 150 template sample. The 44 participants from the first sample were used as atlas inputs, and we again calculated DSC and ICC to compare the manual and automated segmentation approaches.

After validation, we moved on to perform automated multi-atlas based segmentation of the hypothalamus in another sample of participants from our cohort with complete information on primary covariates, laboratory parameters, diffusion-weighted MRI etc. (*n*_2_ = 236, see **Fig. 1**). Again, the 44 participants were used as atlas inputs.

We extracted mean MD from the automatically segmented hypothalami and repeated the multiple regression analysis with age, sex and BMI as predictors. Likewise, we considered hippocampal MD and third ventricular volume as possible confounders. Additionally, since this sample had complete measures of blood pressure, glucose and insulin, we included HOMA-IR and systolic blood pressure into the regression model.

#### Validation of the multi-atlas fusion segmentation

For both the template and the validation sample, we received low to acceptable ICCs (model 3,1) for the volumetric agreement between automatically segmented and manually segmented hypothalamus (Supplementary Table 2). Therefore, we abstained from using volumetric values from this fully automated segmentation in further analyses. Similar to the inter-rater comparison, the DSC between the automatically and manually segmented hypothalami were good with average values across participants of > 0.8 (Supplementary Table 2).

Regarding the MD, we observed good to excellent ICC between the values based on automatically segmented and manually segmented hypothalamic (Supplementary Table 3). In the validation sample 2 the ICC dropped slightly in the left compared to the right hemisphere but remained in the good range (ICC = 0.87).

## Acknowledgments

The authors would like to thank all participants and staff of the LIFE-Adult study. This work was supported by the Deutsche Forschungsgemeinschaft (DFG, German Research Foundation) Projektnummer 209933838 – SFB 1052 project A01 to MS and AV and Projektnummer WI 3342/3-1 to AVW; by the European Union, the European Regional Development Fund, and the Free State of Saxony within the framework of the Excellence Initiative; by the Leipzig Research Center for Civilization Diseases (LIFE) at the University of Leipzig (projects 713-241202, 14505/2470, 14575/2470, and 100329290).

## Supplementary information

### Details about the multi-atlas fusion segmentation

The algorithm included three main steps: First, the atlas images were non-linearly coregistered to the study-specific template using antsRegistration. The registration included a rigid-body transform, an affine transform and the non-linear ‘SyN’ registration step with four resolution levels. For exact settings of the parameters, see publicly available scripts. With the same command, the target image was non-linearly registered to the study-specific template. In a third step, an additional quick registration between the atlas images and the target image in the template space was performed. The quick registration used the same parameters as the full registration, but it excluded the fourth resolution level (**Fig. 4**).

All transforms were concatenated and applied in a single registration step to the hypothalami of the atlas images using antsApplyTransforms. This step yielded multiple labels of the hypothalamus in the target native space.

To fuse these labels, we applied STEPS (Similarity and Truth Estimation for Propagated Segmentations) implemented in NiftySeg (https://github.com/KCL-BMEIS/NiftySegSTEPS) which generated one multi-atlas based hypothalamic segmentation per target image.

**Supplementary Table 1:** Measures of intra-rater, inter-rater reliability and spatial overlap

|  | Left Hypothalamus |  | Right Hypothalamus |  |
| --- | --- | --- | --- | --- |
|  | <u>ICC<sup>a</sup></u> | <u>DSC</u> | <u>ICC<sup>a</sup></u> | <u>DSC</u> |
| <u>Rater</u> |  |  |  |  |
| Rater 1 | .73 | .93 | .82 | .94 |
| Rater 2 | .95 | .96 | .89 | .96 |
| Rater 1 – Rater 2 | .83 | .88 | .88 | .89 |

*ICC: Intra-class correlation coefficient, DSC: Dice similarity coefficient*
<sup>a</sup> ICC (1,1) was used to assess agreement within each rater, whereas ICC (3,1) was used to assess agreement between both raters

**Supplementary Table 2:** Inter-rater reliability and percentage of overlap for left and right hypothalamic volume between the semi-automated segmentation sample and the two different multi-label fusion segmentation samples.

|  | Left Hypothalamus |  | Right Hypothalamus |  |
| --- | --- | --- | --- | --- |
|  | <u>ICC<sup>a</sup></u> | <u>DSC</u> | <u>ICC<sup>a</sup></u> | <u>DSC</u> |
| <u>Rater</u> |  |  |  |  |
| Validation 1 (atlas) (n = 44) | .62 | .85 | .55 | .86 |
| Validation 2 (n = 24) | .67 | .85 | .73 | .85 |

*ICC: Intra-class correlation coefficient, DSC: Dice similarity coefficient*
<sup>a</sup> ICC (3,1) was used to assess agreement between both approaches

**Supplementary Table 3:** Inter-rater reliability and percentage of overlap for left and right hypothalamic MD between the semi-automated segmentation sample and the two different multi-label fusion segmentation samples.

|  | Left Hypothalamus | Right Hypothalamus |
| --- | --- | --- |
|  | <u>ICC<sup>a</sup></u> | <u>ICC<sup>a</sup></u> |
| <u>Rater</u> |  |  |
| Validation 1 (atlas) (n = 44) | 0.97 | 0.98 |
| Validation 2 (n = 24) | 0.87 | 0.97 |

*ICC: Intra-class correlation coefficient, DSC: Dice similarity coefficient* <sup>a</sup> ICC (3,1) was used to assess agreement between both approaches

**Supplementary Table 4:** Group characteristics of the samples used for multi-atlas fusion segmentation. Data is given as mean ± standard deviation and range (minimum – maximum).

|  | Semi-automated | Study-specific template | Validation 1 (atlas) | Validation 2 |
| --- | --- | --- | --- | --- |
|  | n = 338 | n = 150 | n = 44 | n = 24 |
| Age (years) | 55.0 ± 12.4<br>(21 – 78) | 55.1 ± 12.4<br>(21 – 78) | 56.3 ± 10.8<br>(34 – 76) | 59.2 ± 10.9<br>(40 – 78) |
| Sex (males/females) | 176/162 | 76/74 | 24/20 | 15/9 |
| Rater (1/2/both) | 166/152/20 | 73/69/8 | 21/19/4 | 11/9/4 |
| BMI (kg/m<sup>2</sup>) | 26.4 ± 3.9<br>(18 – 43) | 26.6 ± 3.8<br>(18-37) | 26.8 ± 3.8<br>(20 – 37) | 25.9 ± 2.8<br>(20 – 30) |

*BMI: body mass index*

